# Unraveling the Neural Landscape of Mental Disorders using Double Functional Independent Primitives (dFIPs)

**DOI:** 10.1101/2024.08.01.606076

**Authors:** Najme Soleimani, Armin Iraji, Godfrey Pearlson, Adrian Preda, Vince D. Calhoun

## Abstract

Mental illnesses extract a high personal and societal cost, and thus explorations of the links between mental illness and functional connectivity in the brain are critical. Investigating major mental illnesses, believed to arise from disruptions in sophisticated neural connections, allows us to comprehend how these neural network disruptions may be linked to altered cognition, emotional regulation, and social interactions. Although neuroimaging has opened new avenues to explore neural alterations linked to mental illnesses, the field still requires precise and sensitive methodologies to inspect these neural substrates of various psychological disorders. In this study, we employ a hierarchical methodology to derive double functionally independent primitives (dFIPs) from resting state functional magnetic resonance neuroimaging data (rs-fMRI). These dFIPs encapsulate canonical overlapping patterns of functional network connectivity (FNC) within the brain. Our investigation focuses on the examination of how combinations of these dFIPs relate to different mental disorder diagnoses. The central aim is to unravel the complex patterns of FNC that correspond to the diverse manifestations of mental illnesses. To achieve this objective, we used a large brain imaging dataset from multiple sites, comprising 5805 total individuals diagnosed with schizophrenia (SCZ), autism spectrum disorder (ASD), bipolar disorder (BPD), major depressive disorder (MDD), and controls. The key revelations of our study unveil distinct patterns associated with each mental disorder through the combination of dFIPs. Notably, certain individual dFIPs exhibit disorder-specific characteristics, while others demonstrate commonalities across disorders. This approach offers a novel, data-driven synthesis of intricate neuroimaging data, thereby illuminating the functional changes intertwined with various mental illnesses. Our results show distinct signatures associated with psychiatric disorders, revealing unique connectivity patterns such as heightened cerebellar connectivity in SCZ and sensory domain hyperconnectivity in ASD, both contrasted with reduced cerebellar-subcortical connectivity. Utilizing the dFIP concept, we pinpoint specific functional connections that differentiate healthy controls from individuals with mental illness, underscoring its utility in identifying neurobiological markers. In summary, our findings delineate how dFIPs serve as unique fingerprints for different mental disorders.

## 1. Introduction

Mental disorders pose a significant challenge in modern healthcare, affecting millions worldwide with diverse and often complex manifestations [1], [2]. The interplay of genetic predispositions, environmental factors, and neural processes contributes to the diverse and overlapping manifestations of these disorders [3],[4]. Understanding their underlying neural signatures of these disorders is crucial for effective diagnosis, treatment, and management. Unique individual aspects of brain activity, indexed by functional magnetic resonance imaging (fMRI), can predict subsequent psychological distress [5],[6]. This has led to claims that scanning brains to detect and individual’s “brain fingerprint” [7], and utilizing this measure as a predictor of mental health outcomes could be the future of personalized mental health care [8]. In recent years, advancements in neuroimaging techniques, particularly resting-state fMRI (rs-fMRI), have provided invaluable insights into the functional organization of the brain [9]. Rs-fMRI primarily allows for the exploration of intrinsic brain activity, revealing functional networks that exhibit synchronized fluctuations in activity across different brain regions even in the absence of explicit tasks [10]. This resting-state functional connectivity offers a unique window into the brain’s intrinsic organization and has emerged as a powerful tool in mapping the neural correlates of various mental disorders [11].

In contrast to the widely used seed-based functional connectivity (FC) approach [12], which involves selecting a specific brain region (i.e., the “seed”) and measuring the correlation of its activity with other regions to infer functional connections, independent component analysis (ICA) served as a statistical method that can be used to decompose the complex and computationally expensive fMRI data into a set of independent components to dissect the sophisticated contributions of specific neural networks to the manifestation of mental disorders [13], [14]. The utilization of ICA presents distinct advantages over traditional FC analysis in the investigation of the brain’s functional architecture and its implications for mental disorders [15]. Unlike seed-based FC approach, which typically relies on predefined regions of interest and their correlations [16], ICA offers a data-driven approach. This allows for the identification of both overlapping and non-overlapping functional networks within the brain, thereby capturing a more comprehensive view of its complex organization [15]. Furthermore, ICA enables the separation of regions with multiple functional roles into distinct networks, enhancing the precision and specificity of the analysis [17]. In practical terms, ICA aims to separate the observed fMRI signal into statistically independent components, each corresponding to distinct patterns of brain activity [15]. These components represent spatial maps of brain regions that exhibit synchronized activity, providing insights into the functional connectivity within the brain [18].

Another crucial concept in understanding brain function is functional network connectivity (FNC) which refers to the temporal correlations between different brain networks identified through methods like ICA. By examining FNC, researchers can gain insights into how these networks interact and how these interactions are altered in various mental disorders. For instance, Du et al. [19] reviewed the use of FNC to classify and predict brain disorders, demonstrating that FNC can reveal complex interactions that are not apparent when examining networks in isolation. However, the interpretation of FNC can be challenging due to the overlapping nature of brain signals and the potential for spurious correlations. Therefore, it is crucial to decompose FNC into its constituent independent components to better understand the distinct interaction patterns within the brain. Techniques like ICA are essential for this decomposition, enabling researchers to isolate meaningful connectivity patterns from noise [20], [21]The flexibility and adaptability of ICA make it particularly well-suited for uncovering subtle and previously unrecognized patterns of connectivity that might be crucial in understanding the neural underpinnings of mental illnesses [22].

Previous studies have demonstrated the efficacy of ICA in discerning distinct patterns within functional brain networks. For instance, Calhoun et al. [18], applied ICA to resting-state fMRI data, revealing distinct resting-state networks such as the default mode network and the sensorimotor networks. Similarly, a more recent study [19] reviewed the application of ICA in classifying and predicting brain disorders using functional network connectivity. This study highlighted the power of ICA to disentangle complex fMRI data into independent components corresponding to distinct functional networks. By applying ICA to rs-fMRI data, the study identified intrinsic connectivity networks (ICNs), networks of brain regions showing consistent functional connectivity during rest, critical for understanding brain disorders. It also discussed the advantages of ICA in revealing hidden brain activity structures and its potential for clinical applications despite challenges like variability in results and the need for standardization. For example, Garrity et al. [23], used ICA to identify disrupted resting-state networks in patients with schizophrenia (SCZ), revealing abnormalities in the default mode network and the fronto-parietal networks. Similarly, studies by Greicius et al. [24] and Sheline et al. [25] employed ICA to investigate functional connectivity changes in individuals with major depressive disorder (MDD), highlighting alterations in the default mode network and the limbic networks.

ICA has also been instrumental in elucidating the neural correlates of cognitive deficits observed in various mental disorders. For example, Calhoun et al. [26], applied ICA to fMRI data from schizophrenia patients collected during a working memory task, revealing fronto-parietal network disruptions associated with impaired cognitive performance. Similarly, Anticevic et al. [27] and Menon [28] utilized ICA to investigate the functional connectivity changes underlying cognitive dysfunction in individuals with bipolar disorder (BPD) and autism spectrum disorder (ASD), respectively.

These investigations highlight ICA’s role as a powerful tool for uncovering the nuanced alterations in functional brain networks associated with mental disorders. Schizophrenia, a severe and chronic mental disorder characterized by disruptions in thought processes, perceptions, and emotional responsiveness, presents a complex array of neural alterations that challenge traditional diagnostic and treatment paradigms [29]. Recent neuroimaging studies in schizophrenia employing ICA have unveiled aberrant functional connectivity patterns in schizophrenia, particularly within the default mode network and the fronto-parietal network, shedding light on the neurobiological underpinnings of the disorder [30], [31]. Moreover, major depressive disorder and bipolar disorder are additional serious psychiatric conditions characterized by distinct but overlapping neural signatures [32]. Disruptions in the default mode and the limbic networks have been consistently observed in individuals with MDD, likely reflecting alterations in emotional regulation and self-referential processing [33]. Similarly, studies of BPD have revealed perturbations in various functional networks, including those involved in emotion regulation and self-referential processing, underscoring the heterogeneity and complexity of neural alterations in this disorder [34], [35]. By identifying specific patterns of dysfunction within intrinsic and task-related networks, ICA offers a comprehensive view of the neural underpinnings of psychiatric conditions. This approach holds promise for the development of biomarkers, personalized treatment strategies, and a deeper understanding of the complex interplay between brain function and mental health.

The present study proposes a hierarchical approach, introducing the concept of dFIP. The essential idea is to leverage a fully automated spatial ICA approach, called NeuroMark, to estimate functional network connectivity (FNCs), then to impose a second step to identify maximally independent FNC primitives that covary among subjects. These are then used as features to characterize and compare mental illnesses within a large dataset. By leveraging large-scale brain imaging datasets, we aim to estimate dFIPs and evaluate their potential to disentangle the complex neural signatures associated with a variety of major psychiatric conditions. This novel approach aims to extract complex and hierarchical patterns from functional connectivity data, offering a unique perspective on the intricate neural mechanisms associated with mental disorders such as schizophrenia, autism spectrum disorder, psychotic bipolar disorder, and major depressive disorder. A comprehensive understanding of mental disorders requires not only identifying specific patterns but also elucidating the relationships between these patterns across the diverse spectrum of psychological conditions. It is essential to acknowledge the substantial biological overlap among DSM diagnoses. For instance, recent research by Clementz [35] used an algorithmic approach based on features derived from standard clinical assessment scales to correctly identify pre-defined cross-diagnostic “Biotypes.” These biological subtypes were initially created using purely biological information (e.g. eye movements, EEG, cognition). This research highlighted the biologically interplay between schizophrenia, schizo-affective and psychotic bipolar disorder, revealing instances where individuals may exhibit distinct neurobiological signatures, with some demonstrating greater numbers of features characteristic of different disorders, while others manifest a unique profile. The evaluation of combinations of dFIPs in relation to various mental disorders represents an important contribution to the field, promising to reveal distinctive and disorder-specific signatures that may pave the way for more targeted diagnostic and therapeutic interventions. Importantly, the proposed pipeline can be fully automated in the context of our NeuroMark constrained ICA pipeline.

## 2. Materials and Methods

### 2.1. Participants

We evaluate our framework on resting-state functional magnetic resonance imaging (rs-fMRI) data from seven clinical studies including the Bipolar-Schizophrenia Network on Intermediate Phenotypes (B-SNIP) [36], Center for Biomedical Research Excellence (COBRE) [37], Function Biomedical Informatics Research Network (FBIRN) [38], Maryland Psychiatric Research Center (MPRC), Chinese schizophrenia dataset [6], Autism Brain Imaging Data Exchange (ABIDE) [39], and chinese MDD dataset [40]. Participants with head motion > 3° and > 3 mm were discarded. Furthermore, we included additional quality control evaluation of the spatially normalization of functional data across the entire brain for all subjects by comparing the individual mask with the group mask. Initially, individual masks for each subject were generated by identifying voxels exceeding 90% of the mean intensity of the entire brain. A group mask was then computed based on voxels appearing in over 90% of all subjects. Spatial correlations were subsequently calculated between these masks for each subject, considering the top 10, bottom 10, and entire mask slices. Subjects meeting specific correlation thresholds (0.75 for top 10 slices, 0.55 for bottom 10 slices, and 0.8 for the whole mask) were included for further analysis. This method aids the selection of high-quality fMRI data. These criteria yielded a total of N = 5805 individuals diagnosed with SCZ (n=2615, 1302 patients and 1313 controls), ASD (n=1735, 796 patients and 939 controls), BPD with psychosis (n=870, 294 patients and 576 controls), and MDD (n=585, 278 patients and 307 controls).

Table I provide details regarding the demographic information of these individuals.

**Table I.** Basic demographic information associated with multiple psychiatric datasets.

| Disorders | Categories | Age<br>(mean ± std) | Gender<br>(F/M) | Dataset (sample size) |
| --- | --- | --- | --- | --- |
| **SCZ**<br>(**n=2615**) | Patients<br>(n=1302) | 35.8 ± 12.6 | 487/815 | B-SNIP (429), COBRE (68),<br>FBIRN (151), MPRC (147),<br>754-sample SCZ (507) |
| | Control<br>(n=1313) | 34.2 ± 11.7 | 679/634 | B-SNIP (576), COBRE (89),<br>FBIRN (160), MPRC (241),<br>754-sample SCZ (247) |
| **ASD**<br>(**n=1735**) | Patients<br>(n=796) | 16.7 ± 9.2 | 110/686 | ABIDE (796) |
| | Control<br>(n=939) | 16.2 ± 8.6 | 229/710 | ABIDE (939) |
| **Psychotic**<br>**BPD**<br>(**n=870**) | Patients<br>(n=294) | 35.4 ± 12.1 | 192/102 | B-SNIP (294) |
| | Control<br>(n=576) | 36.2 ± 12.5 | 341/235 | B-SNIP (576) |
| **MDD**<br>(**n=585**) | Patients<br>(n=278) | 33.4 ± 11.2 | 169/109 | 585-sample MDD (278) |
| | Control<br>(n=307) | 31.3 ± 10.4 | 191/116 | 585-sample MDD (307) |

### 2.2. Neuroimaging Data and Preprocessing

Rs-fMRI data were preprocessed using standard procedures. FMRI data were preprocessed using statistical parametric mapping (SPM12, http://www.fil.ion.ucl.ac.uk/spm/) under MATLAB 2022 environment. We first discarded the first ten scans. A rigid body motion correction was performed using the toolbox in SPM to correct subject head motion, followed by the slice-timing correction to account for timing difference in slice acquisition. The fMRI data were subsequently warped into the standard Montreal Neurological Institute (MNI) space using an echo planar imaging (EPI) template and were slightly resampled to 3 × 3 ×3 mm3 isotropic voxels. The resampled fMRI images were finally smoothed using a Gaussian kernel with a full width at half maximum (FWHM) = 6 mm. Four additional post-processing steps were performed to further reduce artifacts/noise in the time courses: (1) detrending to eliminate linear, quadratic, and cubic trends; (2) de-spiking temporal outliers; (3) low pass filtering with a cut-off frequency of 0.15HZ; (4) regressing out six head motion parameters and their temporal derivatives. To mitigate and remove confounds, the biological (i.e., age and gender) and technical (i.e., scanner and site) covariates were regressed out for each disorder respectively.

### 2.3. Spatially Constrained Independent Component Analysis

ICA is a widely used data-driven method capable of recovering a set of maximally independent sources from multivariate data [41]. However, one of the challenges that is found in the standard ICA is “order ambiguity”, which indicates that the order of the independent components (ICs) estimated by the standard ICA is arbitrary [42]. While group ICA allows for correspondence across subjects, it may not be fully adaptive to or independent from individual subjects. Additional prior information can contribute to the solution to this problem. Spatially constrained independent component analysis is an advanced technique within the realm of neuroimaging research, specifically designed to enhance the precision of ICA. It operates on the fundamental principle of decomposing fMRI data into spatially distinct and functionally meaningful components, while also incorporating prior (independently derived) spatial information to guide the decomposition process [42]. By imposing spatial constraints derived from anatomical or functional atlases, spatially constrained ICA aims to refine the separation of brain networks and regions of interest, improving the interpretability of the results [20]. This approach allows us to focus on specific brain areas or networks of interest, enhancing the sensitivity to detect subtle changes in functional connectivity associated with various cognitive tasks or clinical conditions.

#### 2.3.1. NeuroMark Pipeline

To capture reliable intrinsic connectivity networks (ICNs) and their corresponding time courses (TCs) for each fMRI scan, the Neuromark_fMRI_1.0 template [43] was applied to the data as implemented in the GIFT toolbox (http://trendscenter.org/software/gift) and also available for direct download (http://trendscenter.org/data), resulting in 53 ICNs. The NeuroMark framework leverages spatially constrained ICA and automates the estimation of reproducible functional brain markers across subjects, datasets, and studies. Unlike methods that focus on fixed brain regions across subjects (i.e., ROI-based approaches), NeuroMark can identify brain networks that remain comparable across individuals while also accommodating the variability present within each person’s networks. Previous research has provided evidence of NeuroMark’s effectiveness in recognizing a range of brain markers and abnormalities within different populations [44], [45]. Moreover, these studies have demonstrated consistent ability of NeuroMark to detect patterns of group differences accurately and to maintain individual classification accuracy over relatively short periods of time [46]. The seven subcategories into which the ICNs are partitioned include subcortical (SC), auditory (AUD), visual (VIS), sensorimotor (SM), cognitive control (CC), default mode network (DMN) and cerebellar (CB) components.

### 2.4. Double Functionally Independent Primitives (dFIP)

The methodological framework for extracting dFIPs employed a hierarchical approach, commencing with the extraction of a total of 53 ICNs for each subject using the NeuroMark fMRI 1.0 template. Subsequently, these ICNs were utilized to compute individual 53 × 53 FNC matrices for each subject defined based on the correlation among ICNs. Next, references for each disorder and its corresponding healthy control group were established by averaging the FNCs of patients and corresponding healthy controls within their respective clinical cohorts. Following this, a second ICA was applied to the vectorized subject-by-1378 FNC matrices of 5805 individuals, resulting in the extraction of 15 component FNCs which constitute the dFIPs, along with their associated component weights. An exhaustive analysis has been done to determine the optimal number of components which involved systematically sweeping through a range from 5 to 60 components, incrementing by 1 at each step, revealing the significant impact of varying component numbers on the outcome. The workflow of the study is illustrated in Figure 1.

**Figure 1.**
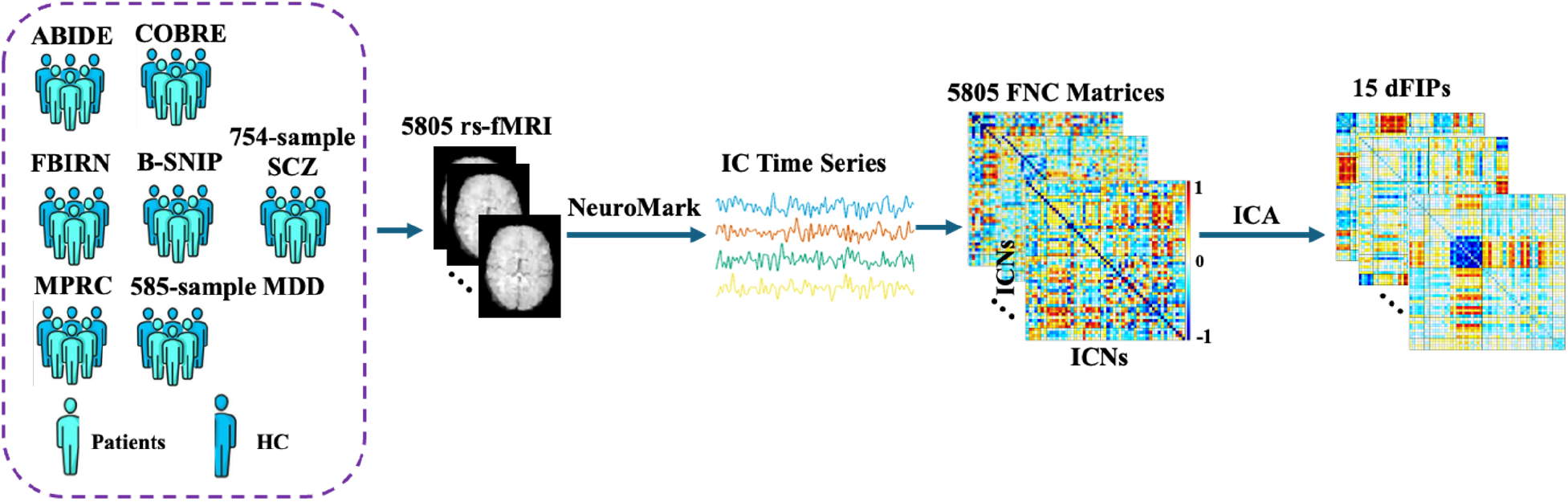
Workflow of the study. The hierarchical approach to extract dFIPs involved first running the spatially constrained ICA pipeline. The NeuroMark fMRI 1.0 template was used to extract 53 intrinsic connectivity networks (ICNs) which was used to compute the 53 × 53 FNC for each subject, followed by employing ICA on all the FNC matrices associated with 5805 individuals to extract 15 double functionally independent primitives (dFIPs).

Following the preprocessing, we performed a two-sample t-test on each of the dFIPs to ascertain significant differences in component loadings between each psychiatric disorder and its corresponding healthy control group.

Next, we conducted multiple regression analyses at both individual and group levels, using each individual dFIP for each clinical group. Following standard regression principles, we employed a comprehensive approach to understand the relationships between distinct variables. At the group level, we used the computed reference as the response variable to explore its intricate connections with each of the 15 dFIPs. Within our analysis, we sought to ascertain the singular contributions of individual dFIPs towards the accurate estimation of this reference, thereby elucidating their respective significance within the broader functional context. At the individual level, the response variable was the subject-specific FNC. Here, we aimed to uncover how each subject’s FNC profile relates to their contributions to the broader functional landscape. This was done to estimate canonical functional patterns that manifest in a given psychiatric disorder. We also employed a method to generate disorder-specific FNC patterns by applying a threshold criterion to the *p-values* associated with beta coefficients. Specifically, we consider only beta coefficients whose −*log*_10_*(pvalue)*is higher than 2 standard deviations (std) from each disorder. This approach serves to isolate and highlight the patterns that exhibit the most pronounced discriminative potential between the respective groups.

To visually portray the discriminative power of each of the 15 dFIPs in distinguishing patients from healthy controls, 2D histograms were generated such that we computed the maximum and minimum values for each component, extracting these indices across both patients and healthy controls. Subsequently, we calculated the difference between these values and plotted the associated 2D histogram. This enabled us to visualize and discern which components exhibited the most discriminative patterns in distinguishing between the two groups, as well as enabling us to visualize the distribution of FNC values for each dFIP. Additionally, equivalent 1D histograms were constructed from the joint 2D distribution to represent marginal distributions. These comprehensive visualizations facilitated a thorough comparison of the impact of individual dFIPs across the spectrum of health and disorder.

The overarching goal of our approach was to identify the specific contributions of each dFIP to the observed intergroup variations. By doing so, it aimed to enhance our understanding of the intricate relationships between functional connectivity patterns and the diverse array of pathological conditions encountered in psychiatric disorders. Moreover, this approach allowed for the identification of connectivity patterns that contribute significantly to the manifestation of psychiatric conditions. Furthermore, it provides a robust and systematic framework for investigating the landscape of functional connectivity in psychiatric disorders. By leveraging robust and automated data-driven pipelines, while also generating dFIPs as a type of ‘FNC basis set’ then using these to derive disorder specific patterns, we hope to provide a way to help unravel the complex interplay of brain networks underlying these conditions.

## 3. Results

### 3.1. Distinct Connectivity Signatures from dFIP Analysis in Psychiatric Disorders

As shown in Figure 2, the FNC references for each mental disorder, after removing covariates, is depicted in the lower triangle, and the respective healthy controls are depicted in the upper triangle. Schizophrenia patients (SCZ) exhibit a heightened connectivity pattern (hyperconnectivity) between the cerebellum and the sensorimotor, auditory, and visual networks, alongside reduced connectivity between the cerebellum and subcortical regions. Similarly, the ASD and BPD group also demonstrated decreased connectivity between the cerebellum and subcortical networks. Individuals with BPD exhibit modest alterations in functional connectivity patterns compared to healthy counterparts, with disruptions in the connectivity between various brain regions including CB and DMN highlighting the alterations.

**Figure 2.**
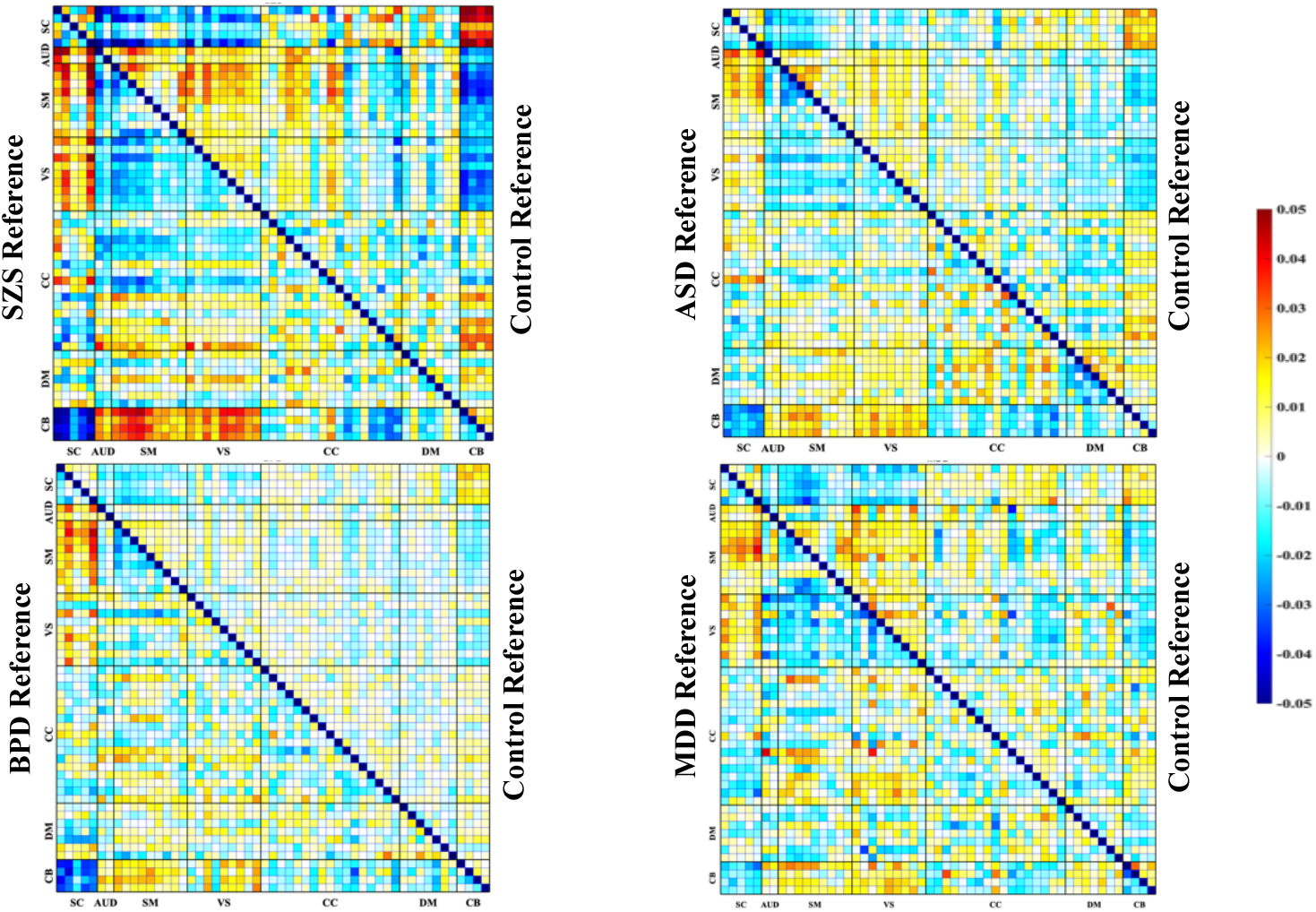
FNC references generated for each psychiatric disorder. The references were obtained for a given disorder and its corresponding healthy control group by averaging the FNCs of patients and controls respectively within each clinical group relative to the overall mean FNC. Each figure displays the FNC of a psychiatric disorder in the lower triangle, and the FNC of the control group in the upper triangle.

The radar plot serves as a graphical representation of the significance −*log*_10_*(pvalue)*of beta weights obtained from the regression analysis performed on dFIPs, offering a condensed depiction of key features reflective of distinct clinical disorder profiles (Figure 3). Despite observable commonalities in dFIP patterns across various disorders, characterized by shared patterns, there also exists notable specificity in these patterns. For example, a dFIP pattern might display weakness within one group while demonstrating strength within others, helping characterize the neural connectivity profiles in different psychiatric conditions. For instance, the impact of dFIP 14 on SCZ and ASD, contrasted with its reduced significance in BPD and MDD, underscores its role as a distinguishing factor between these psychiatric conditions. Upon closer examination of patients diagnosed with SCZ and ASD, who exhibit some degree of overlap in their connectivity patterns, it becomes apparent that certain dFIP patterns play a pivotal role in distinguishing these disorders. Specifically, patterns indicative of hyperconnectivity between the AUD, VIS, and CB domains, coupled with hypoconnectivity between the SC and cerebellar networks, emerged as prominent factors in characterizing the reference profiles for SZ and ASD. Additionally, associations were noted between the DMN and the CC domain in the context of ASD patients, further enriching our understanding of the neurobiological underpinnings of these conditions.

**Figure 3.**
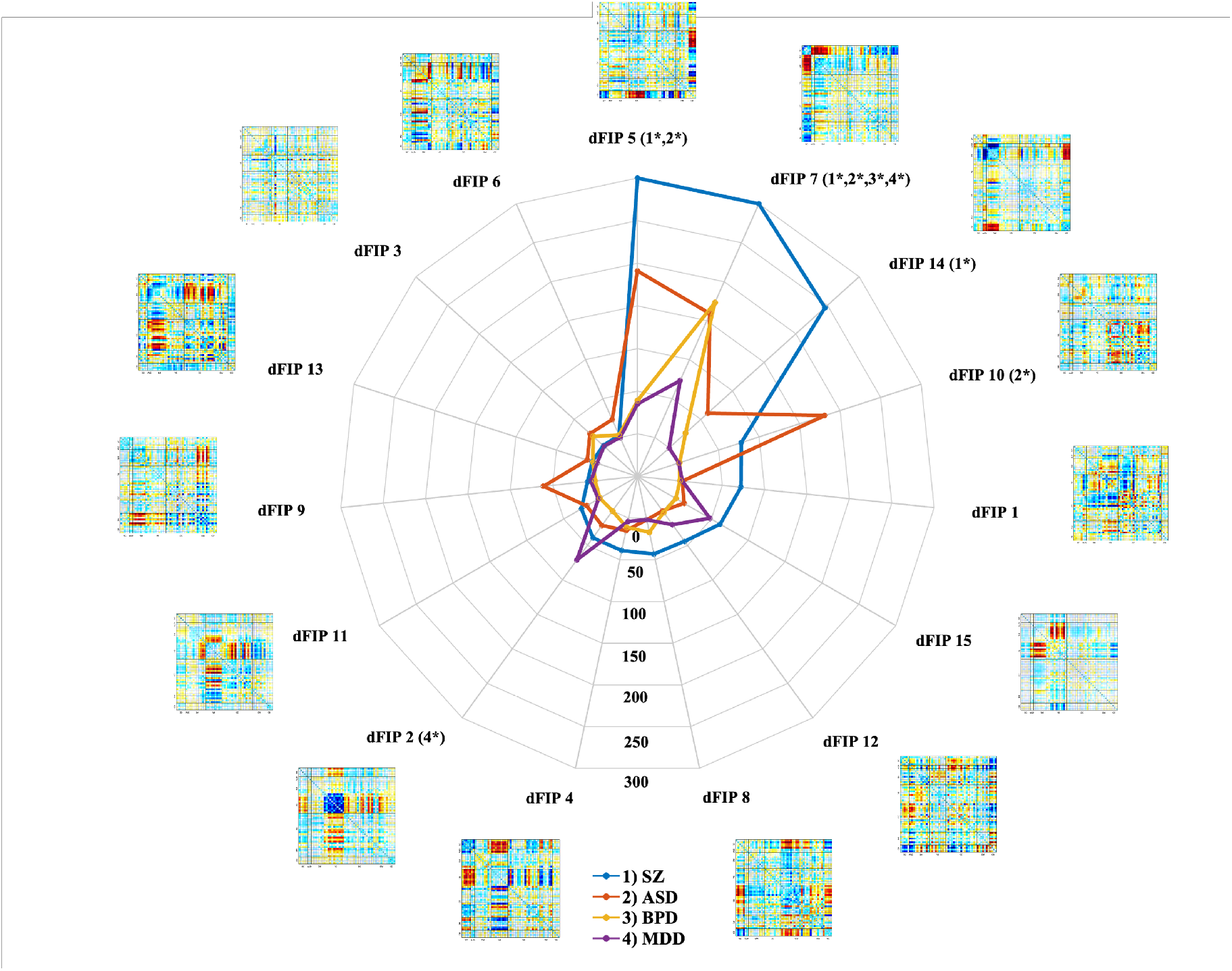
Radar plot associated with the −*log*_10_*(pvalue)*(sorted by the SZ significance from maximum to minimum) revealing the components which play a significant role in elucidating and characterizing the distinct patterns of each disorder. Those components with −*log*_10_*(pvalue)*> 2 *SD* are specified with an asterisk for each disorder.

In contrast, the differences observed in BPD and MDD appear more subtle yet distinct, with specific components assuming significance in the estimation of each disorder. Notably, the seventh component FNC emerged as a key indicator for BPD, while components 2, 7, and 15 exhibited significant importance in delineating patterns associated with MDD. These findings underscore the unique signatures associated with each clinically defined disorder, shedding light on the complex interplay of brain networks implicated in psychiatric conditions.

The incorporation of neurobiological markers into the development of risk assessment tools represents a pivotal advancement in the realm of precision medicine for mental health. By discerning the intricate connectivity patterns underlying several serious psychiatric disorders, these tools hold the potential to enhance diagnostic accuracy and facilitate tailored interventions for individuals at risk. This integrative approach not only deepens our understanding of the neurobiological substrates of mental illness but also lays the groundwork for eventually improving clinical outcomes and optimizing therapeutic strategies.

As illustrated in Figure 4, the reconstructed FNCs offer valuable insights into the distinctive connectivity profiles associated with each clinical disorder, derived from the most salient dFIPs that effectively distinguish patients from healthy controls (HCs). Notably, individuals diagnosed with schizophrenia (SCZ) exhibited significantly heightened connectivity between the AUD, SM, and SC domains, in contrast to the diminished connectivity within the CB regions. A similar yet discernibly weaker pattern was observed in the CB domain among ASD patients, while simultaneously demonstrating heightened connectivity within certain areas of the CC and DMN.

**Figure 4.**
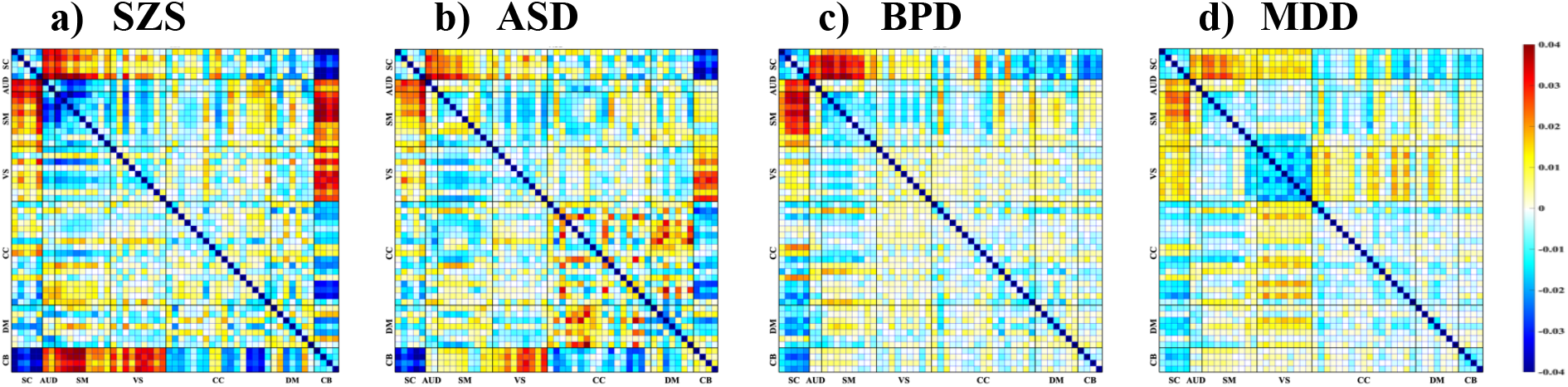
Disorder-specific FNC based on the most discriminative dFIPs whose −*log*_10_*(pvalue)*> 2 *SD* for a) schizophrenia, b) autism spectrum disorder, c) bipolar disorder, and d) major depressive disorder. The construction of disorder-specific FNC profiles reveals the discernible influence of select dFIPs in discriminating between psychiatric conditions.

Conversely, BPD patients displayed subtle yet discernible alterations in their functional connectivity patterns compared to healthy counterparts. Noteworthy disruptions were observed in the connectivity profiles between various brain regions, particularly implicating the CB and DMN domains. Within the MDD cohort, a distinct pattern emerged revealing hypoconnectivity within the VIS domain, accompanied by hyperconnectivity in select regions of the CC domain and DMN when contrasted with the healthy control group. Across SCZ, ASD, BPD, and MDD, commonalities emerge in connectivity patterns, notably within the SM and SC networks. While SCZ and BPD exhibit robust connections between these regions, ASD and MDD demonstrate comparatively weaker connectivity. This shared yet nuanced alteration suggests a potential underlying neurobiological mechanism transcending diagnostic boundaries, implicating the SM-SC (dFIP 7) circuitry in the pathophysiology of these disorders. These regional variations underscore the intricate and diverse neural substrates underlying each psychiatric disorder, presenting a promising avenue for early detection and tailored intervention strategies.

To validate the disorder-specific FNCs as a signature of a particular clinical disorder, we employed a 10-fold cross-validation strategy. In this approach, each iteration utilized nine folds for training purposes while reserving the remaining fold for testing. During the testing phase, the similarity between the test FNCs and the disorder-specific FNCs was quantified using a correlation metric. This analysis demonstrated that the test FNC corresponding to a specific disorder exhibited the highest correlation with the training FNC of that same disorder. Briefly, the diagonal elements of the correlation matrix consistently showed the highest values, indicating that the test FNCs are most similar to the training FNCs of the corresponding disorder (Figure 5). Conversely, off-diagonal elements exhibit relatively lower correlation values, further underscoring the distinctiveness of the FNC patterns for each disorder. For example, the correlation between the SCZ test FNC and ASD training FNC is significantly lower than the correlation between the BPD test FNC and ASD training FNC. This differentiation reinforces the specificity of the FNC signatures to their respective disorders.

**Figure 5.**
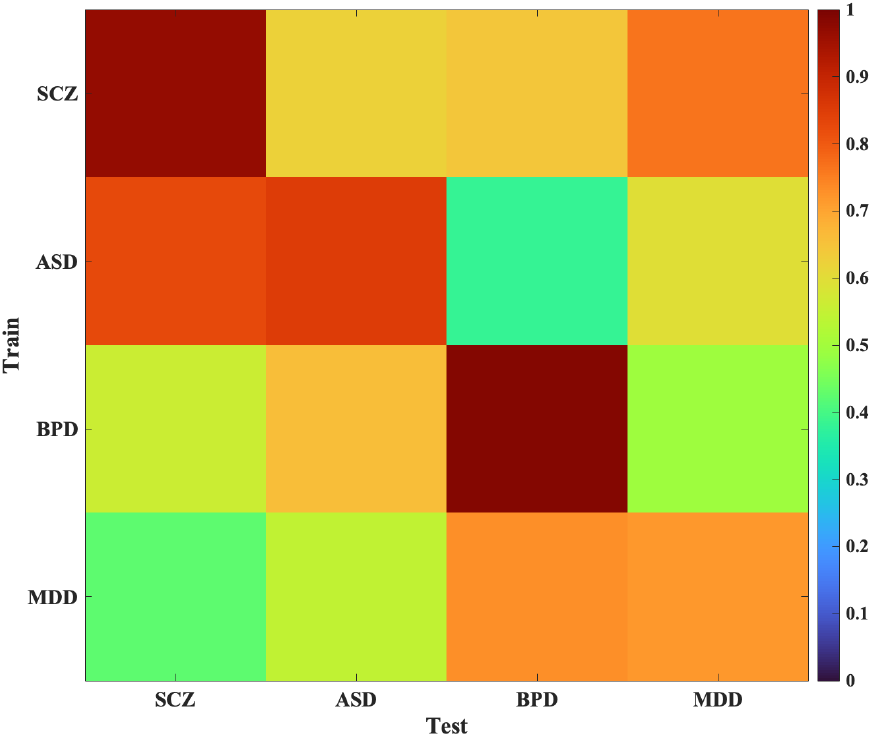
Correlation matrix for disorder-specific FNC validation. This heatmap illustrates the correlation matrix obtained from the 10-fold cross-validation strategy used to validate FNC signatures for four disorders: schizophrenia (SCZ), autism spectrum disorder (ASD), bipolar disorder (BPD), and major depressive disorder (MDD). The rows represent the training FNCs for each disorder, while the columns correspond to the test FNCs. Color intensity indicates the correlation values, with darker colors representing higher correlations. Diagonal elements (e.g., SCZ-SCZ) exhibit the highest correlations, signifying that test FNCs for a specific disorder most closely match the training FNCs of the same disorder.

**Figure 5.**
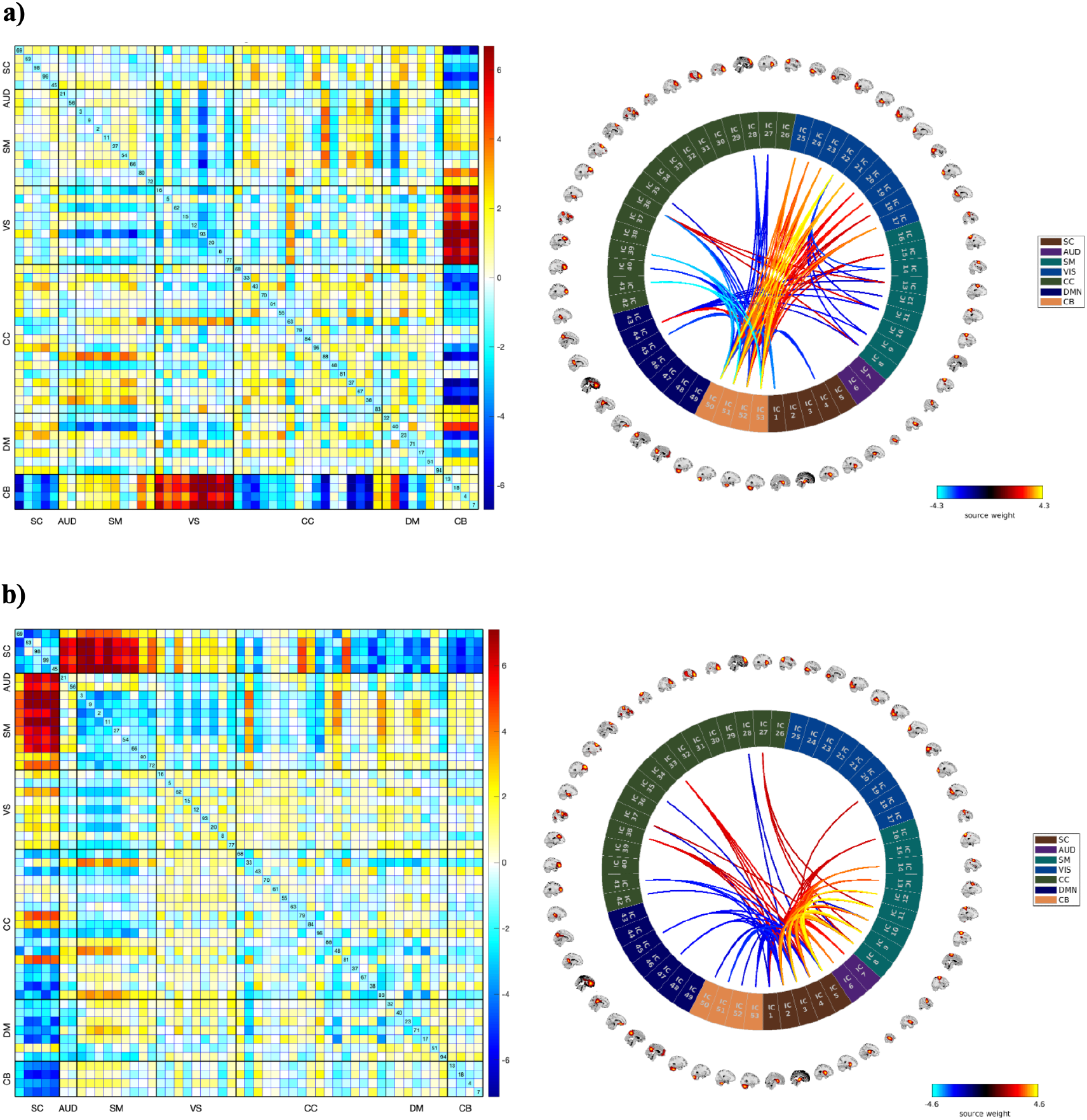
Across various psychiatric disorders, dFIP 5 and 7 emerge as common features, indicating consistent alterations in neural connectivity patterns. a) dFIP 5 and the associated connectogram, and b) dFIP 7 and associated connectogram. Specifically, dFIP 5 exhibits robust connectivity between the CB and VIS domains, alongside weakened connections between the CC and CB regions. This pattern is observed consistently across multiple disorders. Conversely, dFIP 7 demonstrates stronger connectivity between SC and SM regions, suggesting a shared alteration in this neural circuitry among different psychiatric conditions.

### 3.2. Contrasting Signatures of Psychiatric Disorders in Individual Level

A statistical analysis conducted revealed significant differences in the component loadings between patients diagnosed with SCZ versus those diagnosed with ASD when compared to healthy controls. Specifically, individuals with SCZ or with ASD exhibited significant disparities in the loadings across several components, distinct from those observed in the HC group.

More specifically, SCZ patients displayed markedly different loadings on several dFIPs in contrast to the healthy control group, particularly in second, fifth, seventh, fourteenth and fifteenth dFIPs. These dFIPs signify significant alterations in neural connectivity patterns associated with SCZ, highlighting the distinct neurobiological signatures of the disorder. They revealed stronger connectivity between the CB and VIS or SC domains, alongside common and pronounced connectivity between the SM and SC regions. These discrepancies indicate substantial alterations in the functional connectivity patterns within the brain networks associated with SCZ. Notably, the identified differences in component loadings suggest a distinct signature characterizing SCZ, reflecting the unique neural processing and connectivity patterns present in individuals with this disorder.

Similarly, individuals diagnosed with ASD showed significant deviations in the loadings of several components compared to the healthy controls, particularly the fifth, seventh, tenth and fourteenth dFIPs demonstrated significant behavior in AS. These differences signify pronounced alterations in the functional connectivity patterns associated with ASD, highlighting distinct neural network configurations present in individuals with ASD.

In contrast, patients diagnosed with BPD and MDD displayed more nuanced differences in component loadings when compared to the healthy control group, with seventh dFIP emerged as a key indicator for BPD, indicating significant differences in connectivity patterns compared to healthy controls. The alterations observed in BPD and MDD patients suggest subtler changes in functional connectivity patterns, reflecting potential shared neural characteristics within these disorders. Notably, the common pattern observed in SCZ and ASD, characterized by stronger connectivity between SM and CB alongside weaker connectivity between SM and AUD domains (dFIP 14), distinguishes these disorders from BPD and MDD.

Table II provides a comprehensive overview of the statistical results concerning the differences between diagnosis groups (SCZ, ASD, BPD, MDD) and their healthy counterparts in terms of the divergence from the loadings observed in patients compared to healthy controls for each component.

**Table II.** Statistical results associated with various dFIPs in individual level.

| Disorders<br>Components | SCZ |  | ASD |  | BPD |  | MDD |  |
| --- | --- | --- | --- | --- | --- | --- | --- | --- |
|  | *p-value* | *t-value* | *p-value* | *t-value* | *p-value* | *t-value* | *p-value* | *t-value* |
| #1 | 1.6e-08* | 5.66 | 0.36 | 0.91 | 0.90 | 0.12 | 0.35 | 0.93 |
| #2 | 0.001* | 3.25 | 0.04 | 1.99 | 0.90 | -0.12 | 0.007 | 2.70 |
| #3 | 0.19 | -1.28 | 0.004* | 2.87 | 0.08 | 1.70 | 0.42 | -0.79 |
| #4 | 0.0003* | 3.56 | 0.06 | -1.83 | 0.30 | 1.03 | 0.42 | 0.79 |
| #5 | 4.4e-32* | 11.94 | 4.7e-11* | 6.62 | 0.05 | 1.95 | 0.04 | 2.05 |
| #6 | 0.33 | -0.96 | 0.009* | -2.60 | 0.59 | -0.53 | 0.85 | -0.18 |
| #7 | 5.04e-25* | 10.43 | 1.2e-06* | 4.86 | 0.001* | 3.27 | 0.01 | 2.40 |
| #8 | 0.0002* | 3.67 | 0.54 | 0.60 | 0.20 | 1.25 | 0.59 | 0.53 |
| #9 | 0.14 | 1.44 | 0.0004* | 3.53 | 0.90 | 0.11 | 0.37 | -0.88 |
| #10 | 1.3e-08* | 5.69 | 4.6e-13* | 7.29 | 0.71 | 0.35 | 0.77 | -0.28 |
| #11 | 0.002* | -3.06 | 0.03 | -2.16 | 0.84 | 0.19 | 0.47 | 0.72 |
| #12 | 4.6e-05* | -4.08 | 0.92 | -0.08 | 0.69 | 0.39 | 0.025 | -2.24 |
| #13 | 0.23 | -1.20 | 0.10 | 1.60 | 0.56 | 0.57 | 0.84 | 0.19 |
| #14 | 5.2e-27* | 10.88 | 0.0001* | 3.79 | 0.12 | 1.54 | 0.96 | 0.04 |
| #15 | 1.5e-05* | -4.32 | 0.08 | -1.70 | 0.67 | 0.42 | 0.01 | -2.48 |

\* Significant at $p < 0.05$ FDR corrected

To further visualize the neural signatures associated with each mental disorder, we employed a novel approach utilizing FNC extremes across 15 dFIPs. By identifying the maximum and minimum FNC values within the fifth dFIP, we constructed 2D histograms representing the joint distributions of these extremes within our dataset. These histograms were then smoothed using gaussian functions and normalized to reveal probabilities across FNC ranges. Our analysis unveiled distinct marginal distributions between healthy controls (HC) and schizophrenia patients (SCZ). The “dFIP fingerprint” plot, illustrating the difference between HC and SCZ marginal distributions, highlighted regions of significant divergence. Specifically, we observed that the differences in the sums of normalized histograms along the x-axis and y-axis were notably pronounced in SCZ compared to HC. The visualization of the original FNC differences further elucidated patterns of divergence, with positive values indicating higher probabilities in HC and negative values in SCZ (Figure 6). These findings were further supported by the examination of the fifth dFIP values, demonstrating specific functional connections that contribute to the observed divergences. Notably, the differences in FNC patterns could potentially serve as neurobiological markers for schizophrenia.

**Figure 6.**
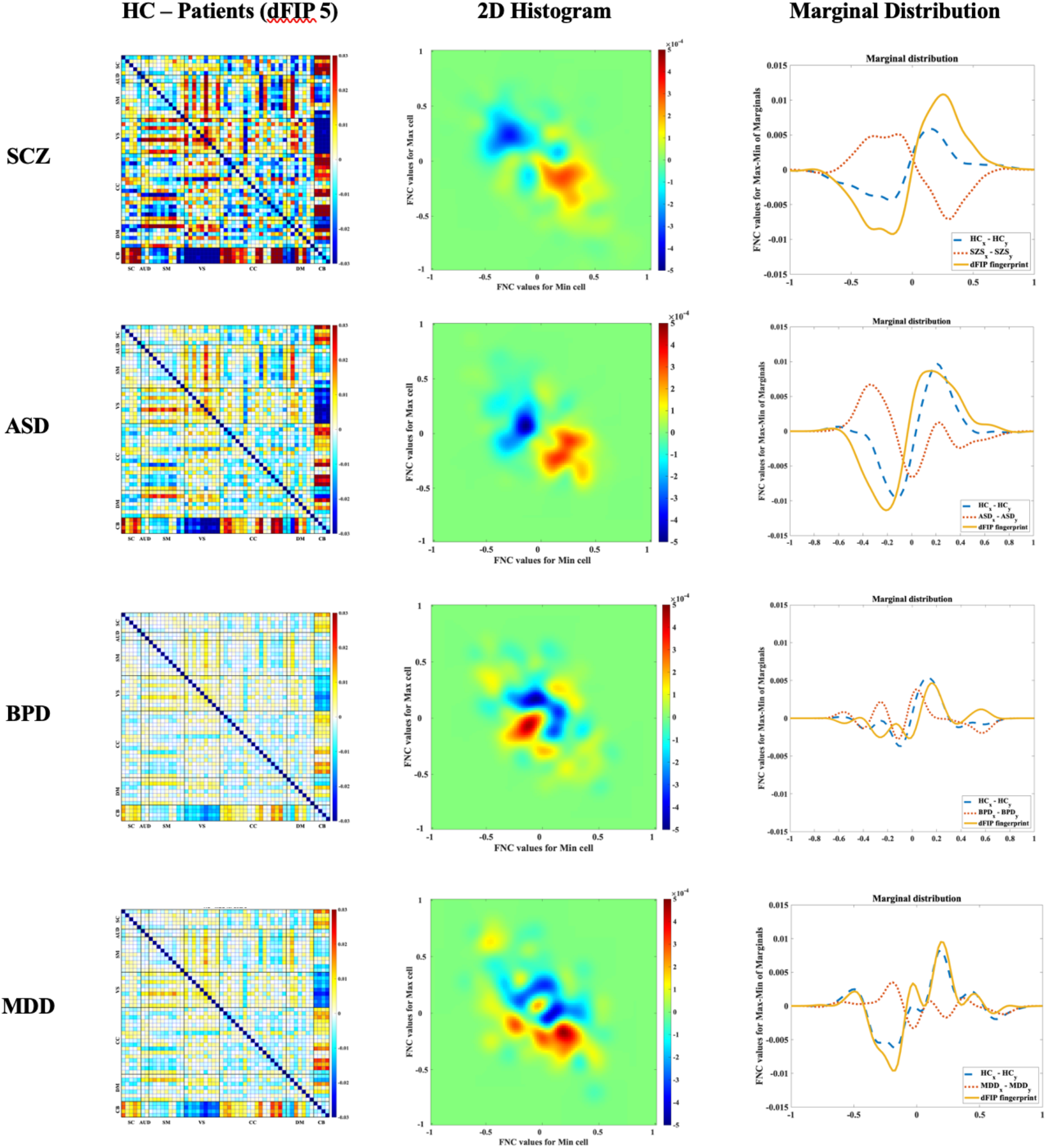
The average HC and patients difference associated with dFIP #5 corresponding with each disorder (first column), the HC – patients in the minimum/maximum joint distribution associated with dFIP #5 for each psychiatric disorder, and marginal distributions for min and max as well as their difference.

### 3.3. Transdiagnostic overlap of mental disorders

Our study further investigated the correlations between the reference patterns associated with each mental disorder to discern overlapping patterns among different disorders. This analysis aimed to shed light on potential comorbidities or shared neurobiological features between conditions. Our findings revealed a notable correlation between the reference patterns of schizophrenia (SCZ) and autism spectrum disorder (ASD), indicating a potential overlap in the underlying neural substrates of these two disorders (Figure 7). This observation underscores the significance of understanding the commonalities and distinctions between mental illnesses, particularly in the context of comorbidity. The identified correlation suggests shared neural mechanisms or pathways that may contribute to both SCZ and ASD, highlighting the complexity of these conditions and the need for integrated approaches in diagnosis and treatment.

**Figure 7.**
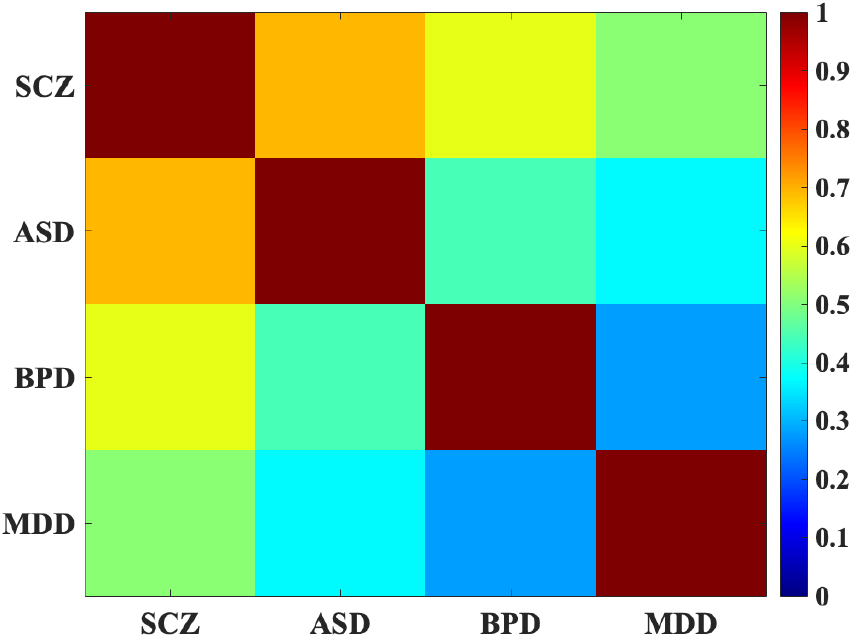
The overlap among different mental disorders. SCZ and ASD exhibit a notable correlation, indicating shared neural mechanisms and potential overlapping neurobiological features between these two disorders.

## 4. Discussion

Understanding the functional connectivity patterns associated with major psychiatric disorders is crucial for advancing diagnosis, treatment, and management strategies. In this study, we employed a novel approach, double functional independent primitives (dFIPs), to unravel the complex neural signatures of schizophrenia (SCZ), autism spectrum disorder (ASD), bipolar disorder (BPD), and major depressive disorder (MDD). Our methodology aimed to provide information about the complex neural alterations linked to these psychiatric conditions, leveraging resting-state functional magnetic resonance imaging (rs-fMRI) data from a large cohort of individuals.

The results of our study revealed distinct neural connectivity signatures for each psychiatric disorder, as evidenced by the differential patterns of dFIPs observed between patients and healthy controls. Notably, patients diagnosed with SCZ exhibited heightened connectivity between brain regions such as the cerebellum and sensorimotor, auditory, and visual networks, along with reduced connectivity between the cerebellum and subcortical regions. These findings align with previous research demonstrating aberrant functional connectivity patterns in SCZ, particularly within the default mode network and the fronto-parietal networks [23], [31], [45]. The observed hyperconnectivity in SCZ patients suggests disruptions in information processing and integration across distinct brain regions, perhaps contributing to the characteristic symptoms of the disorder.

Similarly, individuals diagnosed with ASD and BPD displayed alterations in functional connectivity patterns compared to healthy controls, albeit with some differences. ASD patients exhibited decreased connectivity between the cerebellum and subcortical networks, along with disruptions in the default mode network and cognitive control domains. These findings corroborate previous studies highlighting abnormalities in functional connectivity networks associated with ASD [28], [47]. On the other hand, BPD patients demonstrated subtle yet discernible alterations in connectivity profiles, particularly implicating the cerebellar and default mode network regions. These findings are consistent with prior research indicating disruptions in emotion regulation and self-referential processing networks in BPD [24], [48]

Furthermore, our analysis revealed distinct patterns of functional connectivity associated with MDD, characterized by hypoconnectivity within the visual domain and hyperconnectivity in select regions of the cognitive control and default mode networks. These findings align with previous research demonstrating alterations in default mode and limbic networks in individuals with MDD [24], [48].

The comparison of our results with existing literature highlights the consistency and robustness of our findings, demonstrating the utility of dFIPs in uncovering nuanced patterns of neural connectivity across different psychiatric disorders. Importantly, our study contributes to a deeper understanding of the neural underpinnings of mental illnesses and underscores the potential of sophisticated neuroimaging analyses in psychiatric research.

Moreover, our investigation into the comorbidity of mental disorders revealed correlations between the reference patterns of SCZ and ASD as shown elsewhere [49], suggesting potential shared neural mechanisms or pathways between these conditions. This observation emphasizes the importance of integrated approaches in diagnosis and treatment, considering the complexity and overlapping features of psychiatric disorders. Our findings align with previous research indicating shared genetic and neurobiological factors between SCZ and ASD [50],[51] highlighting the need for comprehensive assessment and personalized interventions for individuals with comorbid conditions.

Overall, our study provides valuable insights into the neural connectivity patterns associated with psychiatric disorders, offering a data-driven synthesis of intricate neuroimaging data. By elucidating the distinct functional network dynamics underlying these conditions, our findings pave the way for the development of biomarkers, personalized treatment strategies, and a deeper understanding of the complex interplay between brain function and mental health.

### Limitations and future work

There are several limitations of the current study that warrant consideration. First, the cross-sectional nature of the data limits our ability to infer causality between the observed neural patterns and mental disorders. Longitudinal studies are needed to determine the directionality of these relationships and to track changes in brain connectivity over time. Second, the use of resting-state fMRI data, while invaluable for capturing intrinsic neural activity, does not provide insight into how these networks function during specific cognitive tasks. Future research could integrate task-based fMRI to explore how these connectivity patterns are modulated by cognitive demands. The variability in fMRI data quality across different sites and scanners also poses a challenge. Even though rigorous preprocessing methods were applied, inherent differences in machine calibration and scanning protocols can introduce biases that might affect the generalizability of the findings. Additionally, the study’s reliance on specific neuroimaging software and statistical methods may limit the applicability of the results across different research settings. In future work we plan to use the dFIPs to study risk in non-clinical sample [2], and potentially to predict future transition to psychosis in clinical high-risk samples [2], as well as looking at heterogeneity including the impact of medication and dimensional measures of psychosis.

## 5. Conclusions

In conclusion, the application of dFIPs represents a methodological advance in the investigation of neural connectivity in psychiatric disorders, offering a comprehensive and nuanced approach to understanding the neural signatures of mental illnesses, by representing disorders as a combination of multiple overlapping patterns of covariation. This study represents a significant advancement in the field of psychiatric neuroimaging by introducing the concept of double functionally independent primitives (dFIPs) to explore the complex signatures associated with various mental disorders. Our findings demonstrate that dFIPs can effectively discriminate between different psychiatric conditions, revealing distinct and nuanced patterns of brain connectivity. These patterns offer potential biomarkers for more precise diagnosis. The use of several large, multi-site datasets enriched the robustness of our findings, showcasing the potential of dFIPs to capture the variability and commonality across different mental disorders. By integrating multiple data-driven techniques, including spatially constrained ICA, this research highlights the intricate interplay of brain networks in conditions including schizophrenia, autism spectrum disorder, psychotic bipolar disorder, and major depressive disorder. Future research should focus on expanding the application of dFIPs in longitudinal studies to track the progression of mental disorders and to validate the predictive power of these neural signatures. Additionally, integrating multimodal imaging data, such as positron emission tomography (PET) or diffusion MRI (dMRI), could provide a more comprehensive view of the neural and molecular mechanisms underlying these conditions. In conclusion, this study health helps advance our understanding of the neural basis of psychiatric disorders and potentially helps move us closer to a more nuanced, personalized approach to mental health diagnostics and intervention. The insights gained from the dFIP analysis promise to help refine our understanding of mental disorders, that will hopefully translate into treatments.

## Acknowledgement

This work was supported by the National Institutes of Health (NIH) grant number R01MH123610 and National Science Foundation (NSF) grant number 2112455 to Vince Calhoun.

